# Bacteria- and temperature-regulated peptides modulate beta-catenin signaling in *Hydra*

**DOI:** 10.1101/747303

**Authors:** J Taubenheim, D Willoweit-Ohl, M Knop, S Franzenburg, J He, TCG Bosch, S Fraune

## Abstract

Animal development has traditionally been viewed as an autonomous process directed by the host genome. But in many animals biotic and abiotic cues, like temperature and bacterial colonizers, provide signals for multiple developmental steps. *Hydra* offers unique features to encode these complex interactions of developmental processes with biotic and abiotic factors. Here, we used the model animal *Hydra* to investigate the impact of bacterial colonizers and temperature on the pattern formation process. In *Hydra*, formation of the head organizer involves the canonical Wnt pathway. Treatment with alsterpaullone (ALP) results in acquiring characteristics of the head organizer in the body column. Intriguingly, germ-free *Hydra* polyps are significantly more sensitive to ALP compared to control polyps. In addition to microbes, β-catenin dependent pattern formation is also affected by temperature. Gene expression analyses led to the identification of two small secreted peptides, named Eco1 and Eco2, being upregulated in the response to both, *Curvibacter* sp, the main bacterial colonizer of *Hydra*, and low temperatures. Loss-of function experiments revealed that Eco peptides are involved in the regulation of pattern formation and have an antagonistic function to Wnt signaling in *Hydra*.

## Introduction

Organisms develop in a specific environment, which is recognized and integrated into developmental programs influencing the phenotype and fitness of an individual. Thereby, environmental factors influencing the developmental processes are very diverse like temperature (1), oxygen levels (2), social interaction (3, 4) or the associated microbiome (5). Adjusting the developmental programs to environmental conditions directly translate into the fitness of an individual and has impact on the evolution (6). The field of Ecological Evolutionary Developmental (Eco-Evo-Devo) Biology integrates these three factors into a theoretical framework (7, 8).

Numerous studies explore the effect of temperature on phenotypic differences and the impact on developmental processes (9–12). However, whether the effect is caused by a mere altered chemical reaction norm or whether temperature is actively sensed and developmental programs are adjusted accordingly, is still under debate. There are evidences for both scenarios (11, 13) and they might not exclude each other.

Similarly, the associated bacteria of an organism have been shown to affect developmental processes of the host (14). They can drive the first cleavage and determine the anterior-posterior orientation of the fertilized egg of nematodes (15), induce the morphogenesis and settlement of tubeworm larva (5), or impact the correct molting event in filarial nematodes (16). In the squid *Euprymna*, bacteria control the development of the ciliated appendages of the light organs (17) and in vertebrates they affect the maturation of the gut (18). How microbial signals or environmental cues are received and integrated into the developmental program of the host is only poorly understood. Sensory nerve cells have been shown to recognize several environmental triggers, like temperature in *C. elegans* (19, 20) and nutrients in *D. melanogaster* (21) and are able to alter phenotypic outcomes during development (22, 23). However, it is unclear whether developmental plasticity has common hubs which are triggered by several environmental cues.

The freshwater polyp *Hydra* harbors a stable microbiota within the glycocalyx of the ectodermal epithelium, which is dominated by a main colonizer *Curvibacter sp*. (24, 25). The microbiota is actively maintained by the host (26–28) and is involved in the protection against fungal infection (25). It appears likely that the microbiota has also an influence on the development of *Hydra*, as constantly occurring developmental processes such as regulation of body size are prone to environmental cues (29). *Hydra* belongs to the phylum of Cnidaria, the sister group of all bilateria. It has a radial symmetric body plan with only one body axis and two blastodermic layers, the endo- and the ectoderm (30, 31). While the stem cells reside in the body column, differentiated cells migrate into the head and foot region (32–35). The constant proliferation and differentiation of stem cells and the migration of cells from the body column into the extremities necessitates ongoing pattern formation processes. In *Hydra*, pattern formation is mainly controlled by a Wnt signaling center in the very tip (hypostome region) of the head (36–38). Transplantation of tissue containing the Wnt organizer can induce a secondary axis in recipient polyps depending on the position of excision and transplantation and follows a morphogenetic field model of diffusion reaction (39–42). The formation of the organizer integrates not only position information, but is also dependent on the surrounding temperature (39). In addition, ectopic activation of Wnt-signaling by the inhibition of the GSK3-β kinase with Alsterpoullone (ALP) is inducing stem cell differentiation and secondary axis formations (43) by translocating and activating the TCF transcription co-factor β-catenin into the nucleus (38). The unlimited stem cell capacity and the constant pattern formation in *Hydra* endows the organism extremely plasticity in terms of regeneration, body size adaption to environment and non-senescence (44–50) On the molecular level temperature acts on the Wnt-TGF-β signaling axis influencing the outcome of developmental decisions such as budding and size regulation of the adult polyp (29). All these processes are reversible indicating a high degree of plasticity of the developmental programs in *Hydra*. How the environmental cues are received and integrated into the developmental program of the animal remains unknown.

Here, we describe the taxonomically restricted gene (TRG) *eco1* and its paralogue *eco2* to be regulated by long-term temperature and microbiota changes in the freshwater polyp *Hydra*. Changes in the expression of *eco* genes adjust the developmental decisions during pattern formation by interference with the Wnt-signaling pathway, controlling axis stability and continuous stem cell differentiation in *Hydra*.

## Results

### Temperature and bacteria modulate beta-catenin activity

To consolidate our previous finding that temperature interferes with the Wnt-dependent developmental program in *Hydra* (29) we treated polyps cultured continuously at 12 °C and 18 °C with Alsterpaullone (ALP) at 18°C (**Figure 1A**). ALP is an inhibitor for GSK3-β and causes an activation of the Wnt-signaling pathway leading to the formation of ectopic tentacles in *Hydra* (29, 43, 51). The number of ectopic tentacles can be used as a proxy to evaluate Wnt signaling strength in the animal, where higher levels of Wnt signaling leads to a higher number of tentacles. Animals reared at 18°C formed ∼40% more ectopic tentacles than animals reared at 12°C (**Figure 1B**), indicating that lower temperatures decreased Wnt-signaling strength and thus plays a role in controlling axis formation and maintenance of the proliferation zone in *Hydra*.

**Figure 1:**
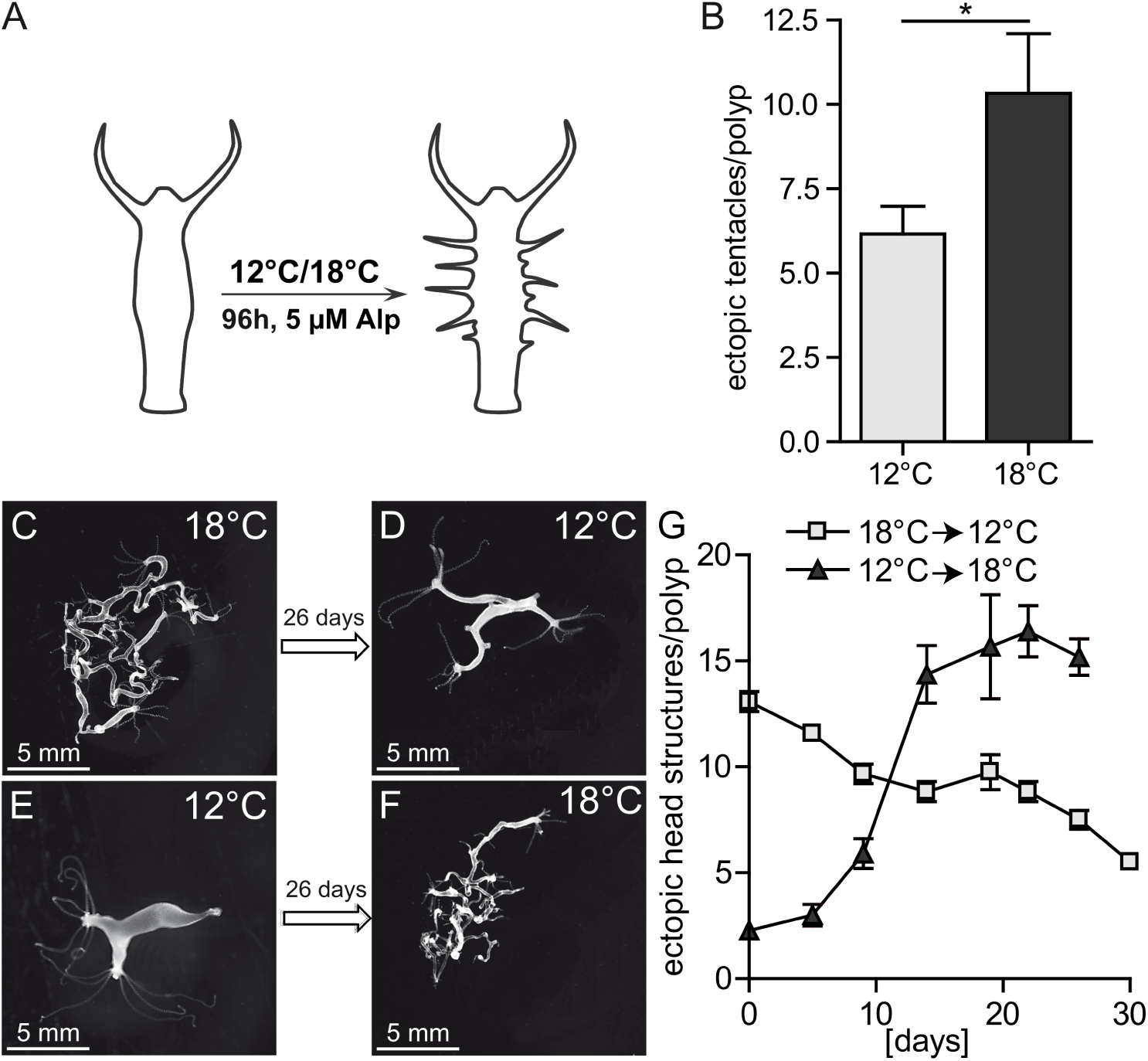
Wnt-signaling is temperature dependent. (A) Animals reared at 12 °C and 18 °C were treated with ALP for 24 h, before assessment of ectopic tentacle formation after 96 h. (B) Lower temperature leads to fewer ectopic tentacle formation after ALP treatment. (t-test, n =12, * p<0.05) (C-F) Constitutive active Wnt signaling in β-catenin OE animals causes multiple heads/body axes at 18 °C, while the severity of the phenotype was subdued when animals were reared at 12 °C. (G) The number of heads produced by the β-catenin OE animals is reversible and depends on the rearing temperature, where surrounding temperatures of 12 °C resulted in fewer heads per polyp (n = 10).

In order to understand how temperature is interfering with the Wnt signaling pathway, we reared transgenic animals, carrying a constitutively active β-catenin over expression (OE) (38) construct at 12°C and 18°C. *Hydra* polyps carrying this construct formed multiple secondary axis and pattern formation is significantly disturbed in these animals (**Figure 1C-F**) (38). Transferring these animals from 18°C to 12°C rescued this phenotype nearly completely (**Figure 1C, D**) by reducing the number of heads in these animals over a course of 26 d (**Figure 1G**). Interestingly, the effect of temperature on axis formation was reversible, as animals with few axis reared at 12°C developed multiple axis within 26 d if cultured at 18°C (**Figure 1E, F, G**). Temperature thereby had neither an effect on the expression of members of the Wnt-signaling pathway nor the β-catenin OE construct (**Supplementary Figure 1**).

To test whether other environmental factors such as the associated microbiota also affects β-catenin dependent development in *Hydra*, we performed the same ALP-treatment on animals with and without associated bacteria (**Figure 2A**). Germfree animals responded nearly four times more to the ectopic activation of Wnt signaling compared to control animals (**Figure 2B-D**). In a second experiment we tested, whether recolonization (conventionalizing) of the germfree animals reduced the increased tentacle formation after ALP treatment and found a significant mitigation of the ALP effect in these animals (**Supplementary Figure 2, Supplementary Table 1**). The observations indicate that not only higher temperatures but also loss of host-associated bacteria increase the Wnt signaling in *Hydra* and affect maintenance of the proliferation zone along the body column.

**Figure 2:**
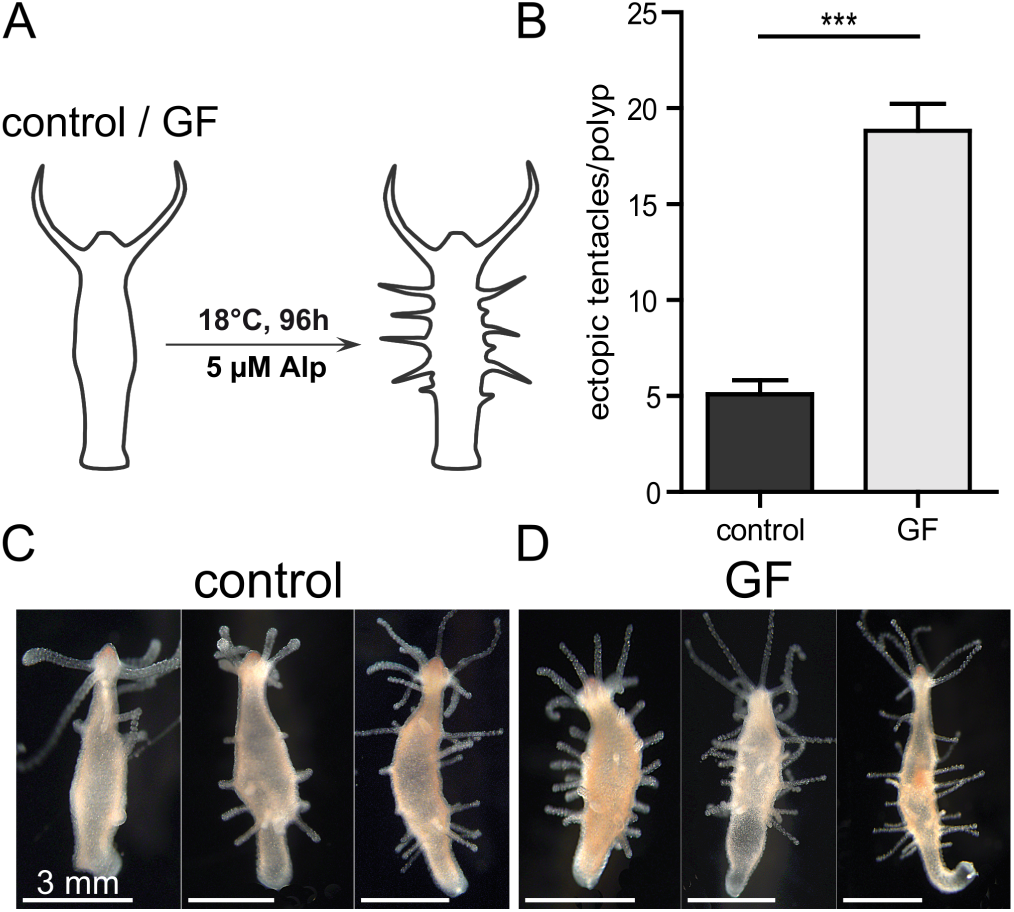
Wnt-signaling is dependent on microbial colonization. (A) Animals were ALP treated for 24 h and number of generated ectopic tentacles were assessed 96 h after treatment, comparing the treatment outcome of germfree (GF) and normally colonized animals (control). (b-d) Ectopic tentacle formation is increased when colonizing bacteria were removed, suggesting a role of microbial colonization in pattern formation in *Hydra* (Mann Whitney U test, n =58, *** p<0.0001).

### Both temperature and bacteria influence the expression of *eco1* and *eco2*

To elucidate the underlying molecular mechanism of the environment-development interaction, we compared differentially regulated genes of a previous microarray study comparing GF to control animals (52) with a recent RNA-seq data set (29), which compared the transcriptome of animals reared at 12°C and 18°C. Both sets of differentially regulated genes overlapped in 55 contigs (**Figure 3A, Supplementary Table 2**). Within this overlap, 18 genes were regulated in the same direction upon lower rearing temperature and removed bacterial colonization (of these 14 [25,45%] down, 4 [7.27%] up). The rest of the genes (37 in total, 67.27%) showed contrary regulation in both conditions (19 [34.55%] down at lower rearing temperature, 18 [32.73%] up at lower rearing temperature) (**Supplementary Table 2**). As low temperature and removal of bacterial colonizers resulted in contrary responses to ALP we searched for genes with contradicting gene expression in this data set. We ranked the genes according their mean expression, fold change and significance level and excluded metabolic genes for further analysis (see methods for details of candidate gene selection). The contig 18166 was ranked highest in this analysis and showed highest differential gene regulation in the temperature experiment and was on position 15 in the bacterial data set among the 55 contigs. Apart from contig 18166 a paralogue contig (14187) was found to be regulated by both temperature and bacteria (position 5 and 18 in the temperature and bacterial data set, respectively). Both paralogues show high sequence homology, encode for small peptides with a predicted signal for secretion, and contain four conserved cysteine residues (**Figure 3B**). For both paralogues no homologs were detectable by BLAST search outside the taxon *Hydra*, indicating that these genes represent taxonomically restricted genes (TRGs) (53, 54).

**Figure 3:**
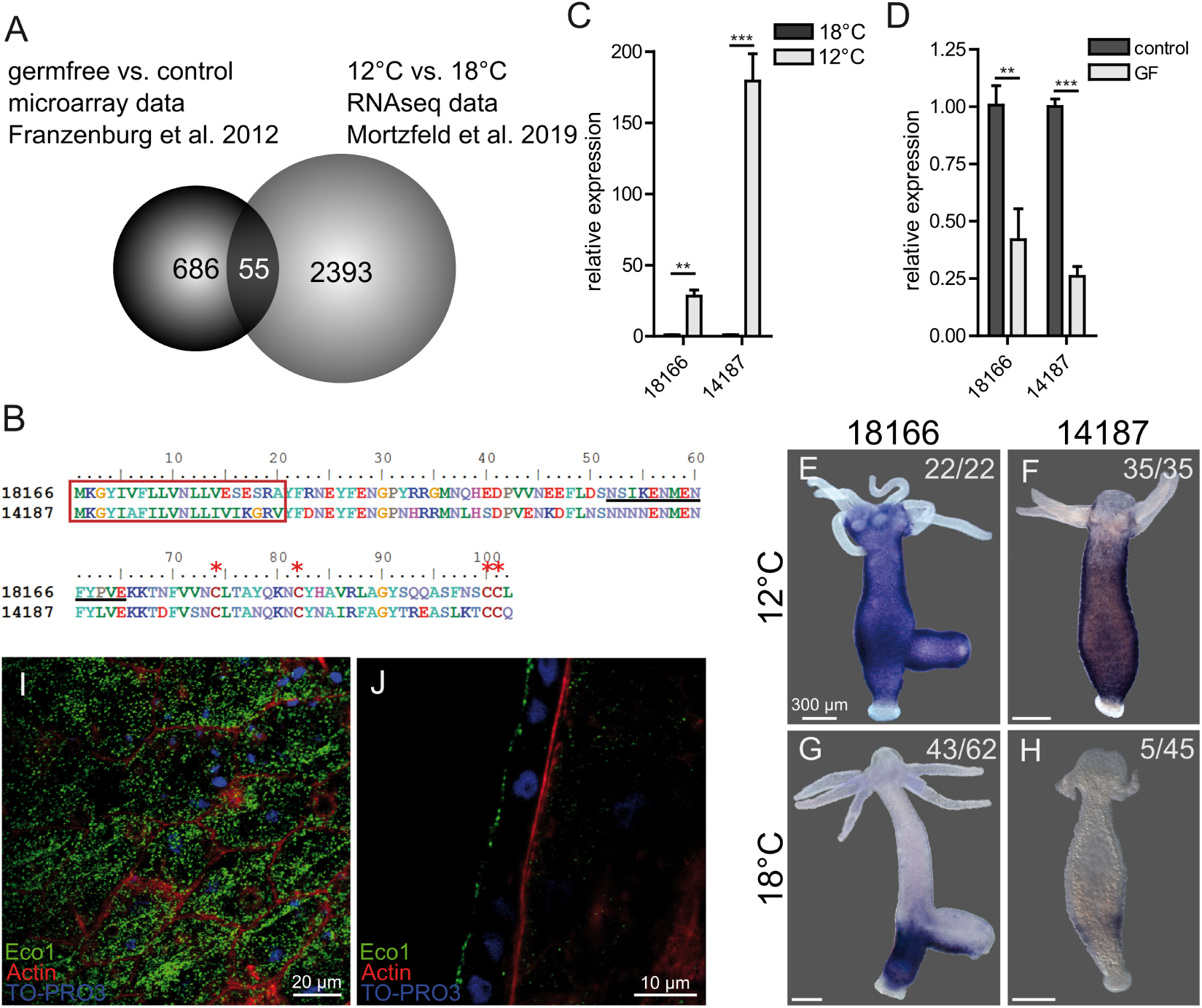
Two TRGs, expressed in the ectoderm, are potential recognition signals for environmental changes. (a) Re-analysis of microarray data, comparing germfree animals with colonized ones, and transcriptomic data, comparing animals at different rearing temperatures, revealed two candidate genes with common regulation upon environmental changes: eco1 and eco2. (b) Sequence analysis of the candidate genes revealed a paralogous relationship between the two genes and a secretion signal peptide, without any domain structure. (c-d) qRT-PCR assessment of eco gene expression in germfree animals and upon temperature shift confirms down regulation of eco1 and eco2 expression at higher rearing temperatures (n =3, Two-way ANOVA, Bonferroni posttests, ** p<0.01, *** p<0.001) and in disturbed microbiome conditions (n =3, Two-way ANOVA, Bonferroni posttests, * p<0.05, ** p<0.01, *** p<0.001). (e-h) eco genes were expressed in the foot and lower third of the animals at 18 °C rearing temperature, but the expression domain expands to the body column and parts of the head at 12 °C. (i-j) Immunohistochemistry with a polyclonal antibody raised against a fragment of Eco1 (underlined in b) show the production of the peptides in the ectodermal epithelium and packaging in vesicles localized in the apical part of the cells, which suggests a secretion of the peptide.

We tested the expression of both paralogues upon temperature (**Figure 3C**) and bacterial cues (**Figure 3D**) via qRT-PCR and observed a significant up-regulation for both genes at low temperature (**Figure 3C**) and in the presence of bacterial colonizers (**Figure 3D**), confirming the initial screening result. Notably, the expression response of both genes to temperature changes was much stronger compared to the bacterial response. The spatial expression pattern of both paralogues were analyzed by whole mount *in-situ* hybridizations. At 12°C, both paralogues are expressed in the ectodermal cell layer along the whole body column, while tentacle and foot tissue showed no expression (**Figure 3E, F**). The expression at 18°C is restricted to the foot in the case of 18166 (**Figure 3G**) and to the lower budding region in the case of 14187 (**Figure 3H**) or showed no detectable level of expression at all (89%) (**Supplementary Figure 3**). In all cases the expression domain of 18166 at 18°C was more expanded than the expression of 14187 (**Figure 3G; H, Supplementary Figure 3**). These results indicate that the expression of both genes is up-regulated due to the expansion of its expression domain from a foot-restricted expression at 18°C to the expression through the whole body column at 12°C.

Using a polyclonal antibody, which was generated against a specific peptide encoded by contig 18166 (**Figure 3B**, underlined sequence), we could observe that the peptide is expressed in the ectodermal epithelial layer and localized in small vesicles (**Figure 3I**) accumulating at the apical side of the epithelial cells (**Figure 3J**). This cellular localization suggest that the peptide is secreted at the apical side of the ectodermal cells. Considering their ecological dependence, we termed the genes *eco1* (contig 18166) and *eco2* (contig 14187), respectively.

### Expression of *eco1* and *eco2* response to environmental changes within two weeks

Having confirmed that both genes respond to changes in temperature and bacterial colonization, we assessed the expression of *eco1* and *eco2* over time in germfree animals and two controls, conventionalized (conv) animals and wildtype polyps (**Figure 4A**). While 8 days post recolonization (dpr) the expression level of *eco1* and *eco2* in conventionalized animals were still equivalent to the levels in germfree (GF) animals, the expression level of both genes recovered within the second week, with rising expression levels similar to control animals (**Figure 4A**). Furthermore, we tested if recolonization with the main colonizer *Curvibacter* sp. alone is sufficient for the regulation of *eco* genes. Analyzing the expression two weeks after recolonization we observed a recovery of the expression levels of both genes (**Figure 4B**), indicating that the specific crosstalk between *Curvibacter* and *Hydra* is sufficient to regulate gene expression of *eco1* and *eco2*.

**Figure 4:**
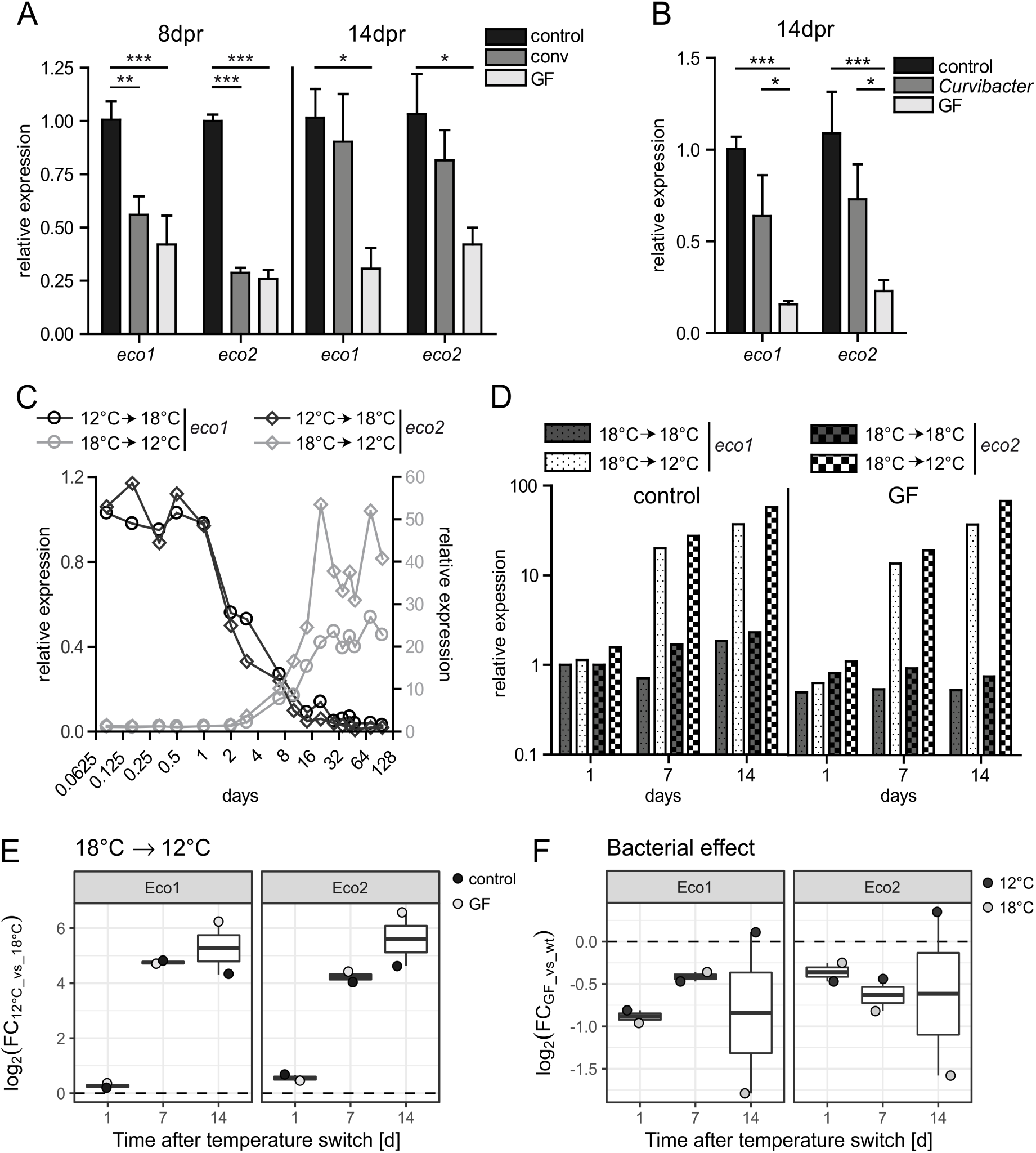
Expression dynamics of eco genes were a long term adaptation to changing microbial state or rearing temperatures. (A) eco gene expression was reestablished by recolonization with bacteria after 14 d, but did not reach normal levels after only 8 d recolonization (n =3, Two-way ANOVA, Bonferroni posttests, * p<0.05, ** p<0.01, *** p<0.001) (B) Recolonization with the main colonizer Curvibacter sp. alone was sufficient to alleviate the expression suppression of eco genes after 14 d of colonization. (n =4, Two-way ANOVA, Bonferroni posttests, * p<0.05, *** p<0.001) (C) eco gene expression changes were established within 14-16 d upon rearing temperature shifts from 18 °C to 12 °C and vice versa. (D) Microbial colonization state and rearing temperature were independent inputs for the expression regulation of eco genes. Gene expression is up regulated upon temperature shifts from 18 °C to 12 °C, independent of colonization state of the animals. (E, F) The graphic displays effect size cleaned of overlaying variances. (E) The expression of *eco* genes increases within 14 days after reduction of rearing temperature, independent of the colonization status of the genes (ANOVA, p = 8.16E-9 for temperature, p = 6.99E-6 for temperature-time interaction, see Supplementary Table 3). (F) Removing bacteria from *Hydra* reduces expression of *eco* genes independent of temperature (ANOVA, p = 0.0557 for colonization, p = 0.20 for colonization-temperature interaction, see Supplementary Table 3).

To get insights into the temporal expression dynamics of *eco1* and *eco2* after temperature change, we transferred animals cultured at 12°C to 18°C and vice versa, and monitored the expression over the course of 28 days (**Figure 4C**). Both genes responded to temperature changes within days, reaching a new stable expression level after around two weeks (**Figure 4C**). Thereby, *eco2* showed a higher up regulation at 12°C compared to *eco1* (**Figure 4C**), which might reflect the fact that *eco2* is expressed at a lower level at 18°C compared to *eco1* (**Figure 3 G, H**).

The fact that both factors, temperature and bacteria, strongly influence the expression of *eco1* and *eco2* raised the question if both factors are interacting with each other. To disentangle both factors, we generated GF animals and maintained them at 18°C or transferred them to 12°C (**Figure 4D-F**). We observed an increase of gene expression in animals transferred to 12°C independent of the microbiota state of the animals (**Figure 4D-E**), while the absence of bacteria reduced gene expression independent of the temperature (**Figure 4F**). The effect of temperature was highly significant in an ANOVA (p = 8.16E-9) which was corrected for gene variation while the bacteria effect was only marginal significant, due to the smaller effect size (p = 0.0557). The interaction term of temperature and colonization was not significant (p = 0.2) indicating no interaction of temperature and colonization in the regulation of *eco* genes (**Supplementary Table 3**). These results suggest independent gene regulation by temperature and microbiota for *eco1* and *eco2*.

In summary, temperature and bacteria dependent regulation of the two genes are reversible and reflect a long-term acclimation to both factors, rather than a short-term regulation and an immediate stress response. The timing of expression changes correlates with the reduction of secondary heads in the β-catenin OE animals reared at 12°C (**Figure 1**). Thus, *eco1* and *eco2* might act as effector genes controlling phenotypic plasticity relaying environmental cues directly to developmental pathways.

### Eco1 and Eco2 act as antagonists to Wnt-signaling

To functionally analyze the role of Eco1 peptides, we designed a hairpin (HP) construct based on the sequence of *eco1* fused to GFP (**Figure 5A**). We generated two transgenic lines (B5 and B8), which displayed constitutive expression of the HP in the ectodermal epithelial cells. The mosaic nature of genetically modified hatchlings allows for the selection of transgenic and non-transgenic lines, which served as genetically identical control lines (except for the hairpin construct). On the level of *in-situ* hybridization the transgenic line B8 showed a dramatically reduced expression level of *eco1* in the whole body column (**Figure 5B**) in comparison to its control line (**Figure 5C**). Checking the knock-down rate of *eco1* by qRT-PCR in both lines revealed a strong down regulation by hairpin-mediated RNAi. Due to high sequence similarity, the hairpin-mediated RNAi targeted also *eco2*, leading to similar down-regulation compared to *eco1* in both transgenic lines (**Figure 5D**). While at 12°C the rate of reduction was between 95%-100% for both genes, the knock-down rate at 18°C was between 20%-40% (**Figure 5D**).

**Figure 5:**
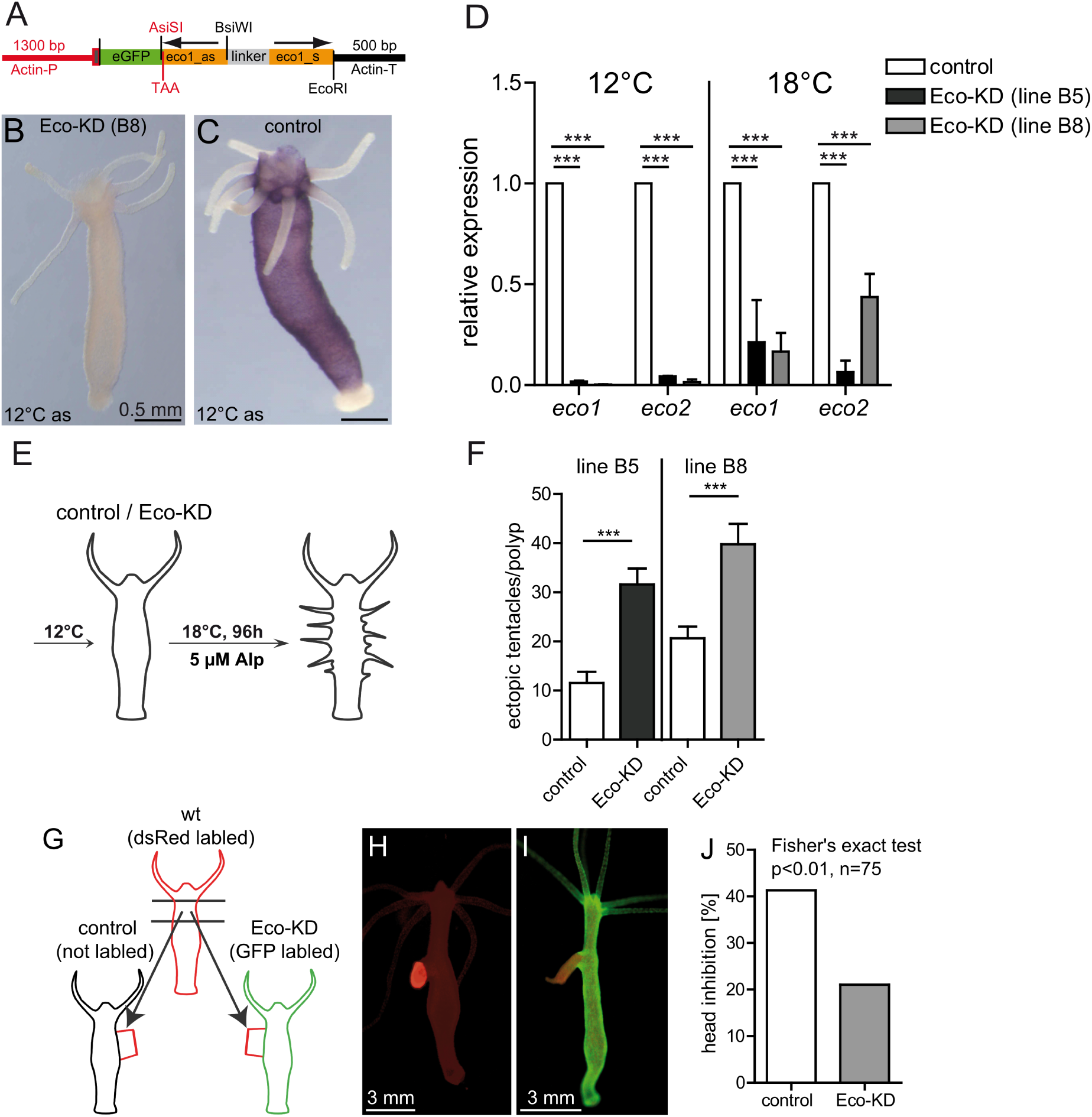
Knockdown of eco genes results in increased Wnt signaling. (a) Eco–hairpin construct for generation of transgenic *Hydra* (as, antisense; s, sense; TAA, stop codon; P, promoter; T, terminator) (b, c) Whole mount in situ hybridization of eco1 expression in Eco-KD (b) and control (c) animals. (d) eco1 and eco2 expression in two transgenic lines (B8 and B5) at 12 °C and 18 °C (n =3, Two-way ANOVA, Bonferroni posttests, *** p<0.001) (e-f) Treating Eco1-KD animals with ALP revealed an increased potential to form ectopic tentacles, indicating a higher Wnt signaling activity. (f) The number of heads formed after transplantation of near head tissue into the body axis of an acceptor polyp serves as a readout for the head inhibition potential, which is governed by the inhibition of the Wnt signaling. (g-h) Control (unlabeled) and Eco-KD (GFP labeled) animals served as acceptor polyps for head tissues from wild type (dsRed labeled) animals to assess the head inhibition potential under disturbed eco expression. (i) Eco-KD polyps showed a reduced head inhibition potential, indicating an impaired Wnt inhibition in these animals. n = 75, p=0.0085, Fisher’s exact test.

To test the hypothesis that Eco peptides act as antagonist to Wnt-signaling we treated both transgenic Eco-knockdown (KD) lines with ALP and compared the number of ectopic tentacles to control animals (**Figure 5E**). We found a significant 3-fold (B5) and 2-fold (B8) increase in number of ectopically formed tentacles in Eco-KD animals, respectively (**Figure 5F**).

Since the Wnt signaling is instructive for head regeneration, we tested, whether Eco peptides are involved in the regeneration process. We cut Eco-KD animals reared at 12°C and 18°C in the middle of the body column, to regenerate a head or a foot (**Supplementary Figure 4A**). During head regeneration we observed no difference neither in the timing (**Supplementary Figure 4B**) nor in the number of tentacles which regenerated between the Eco-KD and control animals (**Supplementary Figure 4C**). Similarly, foot regeneration was not disturbed in Eco-KD animals (**Supplementary Figure 4D**). This result indicates no major role of *eco* genes in the regeneration processes of *Hydra*.

To consolidate the notion of Eco peptides being an antagonist to Wnt-signaling, we performed transplantation experiments to measure the head inhibition potential of Eco-KD animals. In *Hydra*, head activation and inhibition is governed by a gradient of an activator and inhibitor, following a model first described by Alan Turing and specified by Alfred Gierer and Hans Meinhardt (55–57). The model describes a two component system of molecules, which can explain the head forming properties of *Hydra’s* patterning processes. The idea was experimentally tested by Harry MacWilliams in the 1970s and 80s, using transplantation techniques (58, 59). In his experiments he described properties of the head inhibitor and head activator in the animals, using the fact, that head near pieces have organizer functionality (40) and can induce a head in the body column of an acceptor polyp. He described two main findings: first, the rate of head induction increased as the site of transplantation was further away from the head of the donor (head inhibition gradient)(59). Second, the potential to form a head decreases with the distance to the head of the excised pieces (head activation gradient) (58). We used a similar approach and assessed properties of the head inhibition gradient in the Eco-KD background, by comparing the fraction of heads formed in Eco-KD and control animals. To this end, we transplanted head near pieces (directly beneath the tentacle ring) from control donor animals into the body column (approximately one-third from the head) of acceptor animals. We chose the site of transplantation (one-third length from the head) as it was reported that head formation was medium to low in this experimental setting (60). If our notion of decreased head inhibition for the knockdown of the *eco* gene expression was true, we expected an increased fraction of heads formed after transplantation of head near tissue into the body column of Eco-KD animals.

We found, that the fraction forming heads after transplantation is doubled in Eco-KD animals (31 of 75), compared to control animals (16 of 76), indicating a reduced head inhibition potential (**Figure 5G-I**). Together with the ALP experiment, these results demonstrated that Eco peptides antagonize the effects of Wnt-signaling. Lastly, we checked whether *eco* gene expression is regulated by the Wnt pathway and can act as a feedback mechanism in the head formation process. We treated animals with different concentrations of ALP (0.2, 1, and 5 µM) for 24 h and measured *eco* expression via qRT-PCR, but could not detect a regulation of the genes (**Supplementary Figure 5**).

Taken together these results show that the *eco* genes are able to relay environmental signals, like bacterial colonization and temperature, to the Wnt-signaling cascade and by that modulate axis and head formation in *Hydra* according to environmental conditions.

## Discussion

### Wnt-signaling: an evolutionary conserved signaling hub integrating environmental signals to stem cell behavior

The Wnt-signaling pathway most likely evolved in the common ancestor of multicellular animals. Members of the pathway are present in all metazoan animals, but not in fungi, plants, or unicellular eukaryotes, and have regulatory functions in embryogenesis and cell differentiation (61).

In *Hydra* the Wnt pathway is involved in head formation (36), control of bud formation (37, 62) and the differentiation of stem cells (63, 64). Most Wnt family member are expressed in the tip of the hypostome of *Hydra* (37) and the Wnt pathway has been shown to form the activating agent in the head organizer of *Hydra* (36, 37). Wnt corresponds to the head inducer in the Gierer-Meinhardt model, which describes a self-organizing system of head activation (HA) and head inhibition (HI) and is suitable to explain regeneration and axis formation in *Hydra* (42). In this model the HI would correspond to a molecule/gene which is activated by Wnt and is inhibiting the Wnt pathway. Thus, the expression of such a gene would be expected to be expressed in a graded manner from head to foot. Here, we show that Eco peptides antagonizes the Wnt-signaling pathway (**Figure 6**) by reducing both ectopic tentacle formation after ALP treatment (**Figure 5F**) and by transplantation experiments (**Figure 5H-J**). However, the expression domain of *eco* genes is restricted to the foot at 18°C and expands to the head region only, if the animals were reared at 12°C for at least 14 days (**Figure 3E-H, Figure 4C**). The expansion of the expression domain of *eco* genes corresponds to a dramatic increase of the gene expression level, which seems not to appear graded in any form (**Figure 3C, E-H**). Eco peptides do thus not equal the proposed HI of the Gierer-Meinhardt model and therefore add another layer to the Wnt-pathway regulation in *Hydra*.

**Figure 6:**
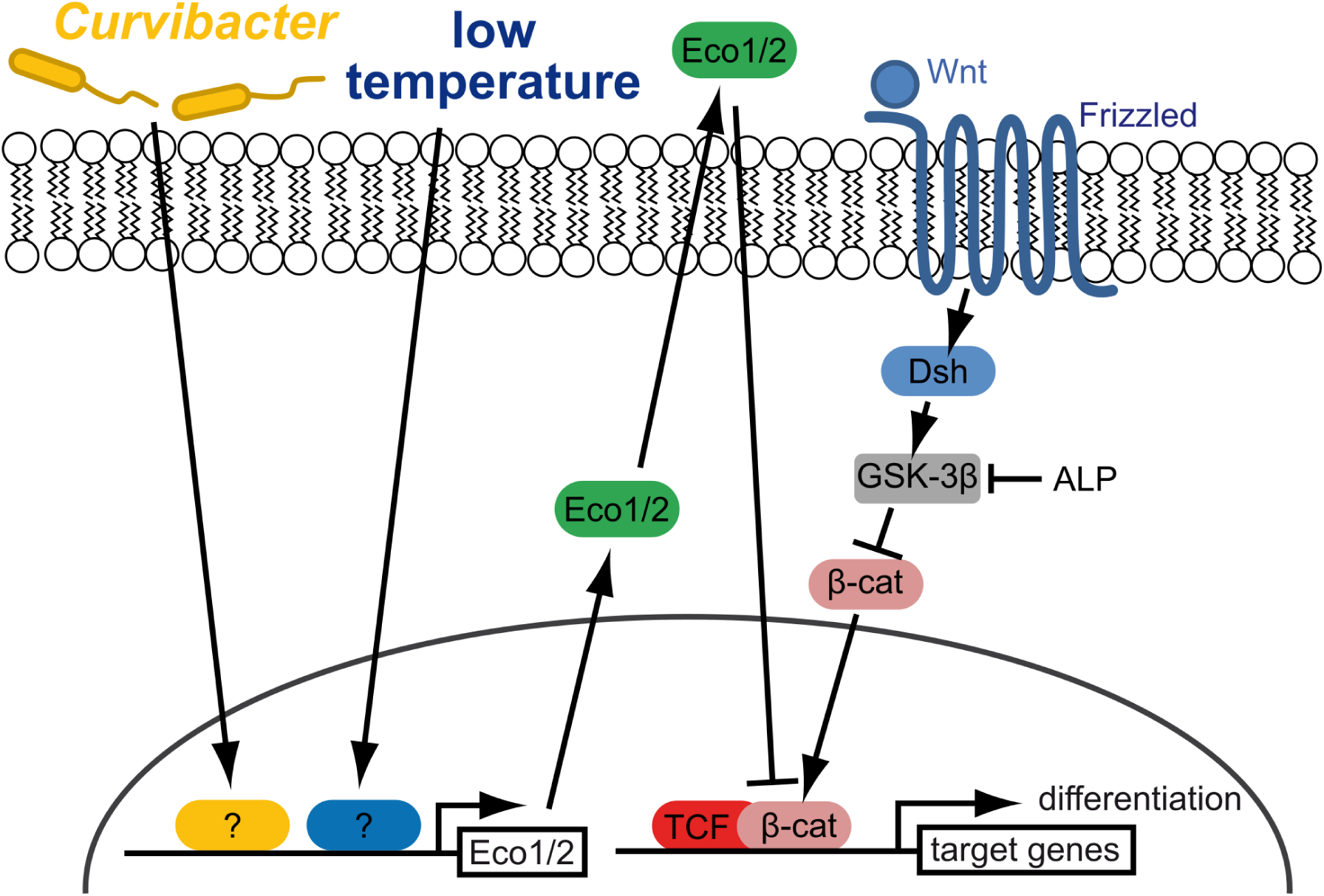
Eco peptides are environmentally triggered inhibitors of Wnt signaling. Microbial colonization and low temperature are independent inputs for Eco peptides and are able to induce its transcription. Eco peptides are synthesized as small peptides, packed into vesicles and secreted into extracellular matrix of the ectoderm. By an unknown mechanism Eco peptides are recognized and the Wnt signaling is inhibited. The inhibition takes place at the level of β-catenin stabilization or the expression of Wnt target genes as Eco expression were able to inhibit activation of the pathway at the GSK-3β level.

The properties of the HI in *Hydra* were elaborately determined by transplantation experiments by MacWilliams (39). Inhibitor level changes occurred during regeneration within 6-24 h after ablation, indicating that the here described inhibitor of the Wnt signaling acts on completely different time scales, than the immediately responding HI. MacWilliams further noted that temperature changes had an influence on the HI potential of *Hydra*. However, these temperature changes were also applied only several hours before transplantation, where *eco* expression changes have not occurred (**Figure 4C**). We therefore argue, that Eco peptides do not contribute to the immediate organizing function of Wnt during head formation, but is rather adjusting background levels of Wnt signals throughout the body. This notion is confirmed by the undisturbed regeneration of Eco-KD animals (**Supplementary Figure 4**) and the otherwise normal appearance of the polyps (including regular budding). A regulation of the background Wnt signaling strength by Eco peptides seems reasonable because Wnt signaling is important for size regulation in *Hydra* by means of bud initiation (29, 62) and body size is regulated with temperature (29, 49). In addition, the variation of multiple axes in the β-catenin OE animals at different rearing temperatures occurred in the same time frame as *eco* gene regulation upon temperature shift (**Figure 1C-G**). Altogether, this could imply a function of Eco peptides in adjustment of Wnt signaling strength as they are regulated with temperature and antagonize Wnt signaling. However, Eco-KD animals showed now obvious size differences in our cultures, which indicates, that the temperature-Wnt regulation seems to be more complex and can not be reduced to Eco signaling alone.

Notably, the proposed HI of the Gierer-Meinhardt model remains enigmatic on the molecular level till today, though various candidates have been proposed, for instance a Hydra dickkopf homolog (65, 66), the transcription factor Sp5 (67), or the glycoprotein thrombospondin (68), but none of them seem to resemble all assumed properties of the HI in the Gierer-Meinhardt model. Further, several downstream genes of the Wnt signaling pathway have been shown to be involved in specific aspects of Wnt signaling, for example the homeobox gene *rax* is important for tentacle formation (69), PKC signaling seems to be important for head regneration (70), but not in bud formation (71), while ERK signaling was essential for both processes (70, 71). We thus argue, that several levels of Wnt-inhibition exist and that the activity of Eco peptides provides one mechanism. Until now, it is unclear how Eco peptides antagonizes Wnt pathway, but we assume that they act downstream of β-catenin given the following arguments. First, we showed that Eco-KD animals have a higher probability of forming tentacles after ALP treatment, but have normal regeneration capability, thus Rax could be a target transcription factor inhibited upon Eco signaling (69). Secondly, we showed that low temperature is antagonizing the effect of over expressed β-catenin. As β-catenin was mutated in a way that it was stabilized against APC dependent degradation (38), we argue that the antagonistic activity is most likely downstream of β-catenin and independent of the APC pathway. Therefore, we predict the presence of an endogenous receptor that binds Eco peptides and activates a signaling cascade antagonizing β-catenin activity.

To date the inputs of *eco* gene regulation on the molecular level are unknown. We can exclude, that Wnt signaling actively regulates *eco* genes, which would be expected if Eco signaling is involved in the immediate head formation process. However, one could speculate on acyl-homoserine-lacton (AHL) as a possible recognition molecule. We previously showed that AHLs produced by *Curvibacter* are quenched by the *Hydra* host to avoid detrimental gene expression of *Curvibacter* on the host (24). This direct interaction indicates an active recognition of AHLs by the host and would mediate a possible route of regulating *eco* gene expression. From the microarray data we can exclude also a MyD88 mediated *eco* regulation, as MyD88-KD animals showed no differences in expression level of *eco* genes (52).

Predicting inputs for the temperature signal is unequally more difficult, as to date no definite temperature sensing mechanism has been found in *Hydra*. TRPs are known to sense high and low temperature and have been identified in *Hydra* (72). TRPM3 has also been linked to high temperature sensation (heat shock) in Hydra (73) but if similar mechanisms exist for cold temperature or whether they can mediate the long term effect of *eco* gene expression is unclear to date. *Eco* expression might be regulated by other head inhibiting substances as MacWilliams has suggested, that physical properties of the HI change at different temperatures, resulting in a longer range of action at lower temperatures, which eventually might translate in an activated *eco* gene expression (39).

Increased Wnt-signaling causes the stem cells in *Hydra* to differentiate and to lose their potential to self-renew (63, 64). Interestingly, the expression of both Eco peptides is regulated by the presence of *Curvibacter*, the main bacterial colonizer of *Hydra*, and low temperature. Individual pathways may signal both external signals to the promotor of *eco* genes, as both factors regulate the expression of both genes independently (**Figure 6, Figure 4D**).

Our model suggests, that the ratio of recognized Wnt and Eco peptides by an individual cell determines the activation of Wnt-target genes. Thereby, the opposing signaling outcomes of Wnt and Eco would determine the degree of differentiation in the tissue, as increased Wnt signaling causes the differentiation of stem cells (63, 64). This process is mediated by Myc1 in interstitial cells, which is directly affected by the increase of Wnt signals due to ALP treatment (64). Given, that Eco peptides are able to counteract increased Wnt signals, we thus argue that the *eco* genes integrate environmental signals into the developmental program and thereby promote the stemness of the cells in the body column of *Hydra*.

There are several studies in vertebrates that showed the integration of environmental signals into the Wnt pathway. In zebrafish, the intestinal bacterium *Aeromonas veronii* enhances β-catenin stability after recognition by MyD88 resulting in higher cell proliferation in the intestine (74). In human epithelial cells the CagA peptide produced by *Helicobacter pylori*, activates β-catenin, leading to transcriptional upregulation of genes implicated in cancer (75, 76). In addition, activation of the aryl hydrocarbon receptor by natural ligands, which are converted from dietary tryptophan and glycosinolates by intestinal microbes (77, 78), results in the degradation of β-catenin (78) and the suppression of intestinal carcinogenesis. We showed another mechanism, how bacterial signals are integrated into the Wnt signaling pathway, via an orphan peptide not present outside the *Hydra* genus. All described mechanisms are highly specific to each of the different model systems (MyD88, CagA, AhR, Eco1) and thus most likely evolved independently, which highlights the necessity of individual adaptation dependent on the life styles to cope with changing environments. However, conserved developmental signaling like the Wnt pathway seems to form signaling hubs to integrate environmental cues. The Wnt signaling pathway thereby fulfills central tasks in pattern formation as well as stem cell regulation in all metazoans and is clearly an important target to mediate developmental decisions upon environmental changes.

### Orphan genes as circuit of environmental cues to host development

This study adds another example to taxonomic restricted or orphan genes as important factors for adaptations and stresses the importance of these fast evolving genes for the interaction with the environment. Orphan genes have been found to play important roles by recruiting new energy resources (79, 80), and occupying new habitats (81). Orphan genes often display functions in developmental programs (81, 82), immunity (26, 83) and mediate interaction with the environment (54, 84, 85). Here we support this notion by showing that the two *eco* paralogues interact with the environment in a way that the current conditions for temperature and microbiota association are sensed and signaled to the Wnt pathway. Thereby, the developmental program is tuned by environmental factors via the regulation of the two orphan peptides Eco1 and Eco2.

### Environment matters

Animal development has traditionally been viewed as an autonomous process directed by the host genome. In recent years, it got evident that biotic and abiotic cues provide a variety of signals that are integrated into the developmental program. These observations resulted in the Eco-Evo-Devo concept (7, 8). Our results provide new evidences for this concept on different levels. Firstly, our study supports the idea that development is plastic and responsive to abiotic (temperature) and biotic (microbiota) factors, as *eco* gene expression is highly dependent on these factors. Secondly, we show that evolution of new traits does not necessarily demand the change in core developmental pathways, but that newly obtained genes can act as modulators of these pathways mediating a gain of function in developmental programs. Thirdly, we show that evolutionary young genes, which modulate conserved developmental signaling cascades, can mediate phenotypic plasticity and increase robustness of patterning formation after disturbance.

## Material and Methods

### Animal culture

All experiments were carried out with either wild type or transgenic *Hydra vulgaris AEP (Hydra carnea* (86, 87)) which were cultured in Hydra medium (HM, 0.28 mM CaCl_2_, 0.33 mM MgSO_4_, 0.5 mM NaHCO_3_ and 0.08 mM KCO_3_) at 18 °C with a 12/12 h light-dark cycle. Animals were fed *ad libitum* 2-3 times the week.

### Transgenic animals

The β-catenin over expression construct contained a truncated β-catenin (AA Δ1-138, containing the phosphorylation site important for Wnt signaling transduction) obtained from *Hydra magnipapillata* terminally fused to eGFP in a modified Hot-G Bluescript II SK + backbone (alias: pHyVec4, Steele, Genebank: DQ385853) (38). The Eco1 hairpin constructs were cloned into a pGEM-T backbone obtained from *Hydra vulgaris* AEP (*Hydra carnea)* sequences (Compagen: HAEP_celera_v1_contig18166) containing base pair 3-303 with a 307 bp linker. dsRED and eGFP labelling constructs were codon usage optimized for *Hydra* and cloned into pGEM-T expression vectors. All constructs were driven by constitutively expressed *Hydra magnipapillata* actin promoters (∼1.5 kb upstream the actin gene) and terminated by an actin terminator (∼1 kb downstream of the gene). Stable transgenic animals were generated by microinjection of genetic constructs into 2-8 cell blastomere of *Hydra*. Transgenic animals hatched with a mosaic expression of eGFP, which served for selection of transgenic cells. The asexual mode of reproduction was used to select for transgenic cells and generate full transgenic animals on the on hand, while on the other hand non transgenic cells were selected to generate a non-transgenic line, which served as a control, containing the same genetic background.

### Antibiotics treatment and recolonization

To render animals germ-free, we treated *Hydra* with a cocktail of ampicillin, rifampicin, streptomycin, and neomycin (50 µg/ml each) in sterile HM for two weeks and daily exchange of the medium. To control for germ-freeness a randomly chosen polyp from each batch were homogenized and plated on R2A-Agar, missing CFU indicated removal of bacteria. Furthermore, whole DNA from a randomly chosen polyp of each batch were isolated (DNeasy Blood & Tissue Kit, *Quiagen*) and a PCR for bacterial DNA contamination was performed (Eub-27F: AGAGTTTGATCCTGGCTCAG, Eub-1492R: GGTTACCTTGTTACGACTT).

To control for the effect of antibiotics in our experiments, animals were conventionalized by recolonizing them with naturally hosted bacteria after antibiotic treatment. To this end, for each germ-free *Hydra* one untreated animal were homognized, resuspended in 100 µl autoclaved HM and added to ∼4 ml HM containing the germ-free animals. The animals were washed 24 h after recolonization with sterile HM to remove excess bacterial load.

### Alsterpaullone experiment

Animals were fed one day prior experiment and treated with 5 µM Alterpaullone (ALP, *Sigma-Aldrich*) for 24 h to inhibit the glycogen synthase kinase 3 β (GSK3-β) and activate the Wnt signaling pathway at 18°C, regardless of previous rearing conditions. After treatment, the polyps were washed and incubated in HM before assessment of ectopic tentacle formation under the binocular 4 d after treatment. To regard any form of the temperature reaction norm we assessed the number of tentacles per polyp after tentacles were clearly developed so that a further increase in number of tentacles could not be observed, even at later time points.

### Temperature treatment of β-catenin animals

β-catenin over expression animals were reared at either 18 °C or 12 °C and standard conditions prior experiment. Animals were transferred to treatment temperature HM (18 °C → 12 °C; 12 °C → 18 °C) kept as single polyps in a 12-well plate. Head structures of the visually largest animal were assessed at day 0, 5, 9, 14, 19, 22, 26, and 30. Smaller animal fragments, which might have appeared during the experiment, were removed from the cavity.

### RNA extraction, cDNA generation, qRT-PCR

Total RNA was isolated by dissolving 15-25 animals in 1 ml TRIzol (*Invitrogen*) by vortexing, after starvation for 2 days. After addition of 300 µL chloroform (p. a.), samples were centrifuged (12,000 g at 4°C), the resulting aqueous phase was transferred in a clean tube, and 400 µl ethanol (99.9 %) was added. The solution was cleaned and desalted using the silica column protocol from Ambion PureLink RNA Mini Kit (*Thermo Scientific*), including an on column DNAse digestion step. 650 ng RNA was used to generate cDNA applying the protocol of the First Strand cDNA Synthesis Kit (*Fermentas*). Resulting cDNA was equilibrated within each experimental setup by semi-quantitative PCR using primer for the house keeping gene ELF1a. Equal amounts (according to PCR quantification) were used with the GoTaq qPCR Master Mix (*Promega*) in a 7300 real-time PCR system (*ABI*). For data analysis the simplified method after Pfaffl (88) was used, assuming a primer efficiency of 100 %.

### Gene selection from transcriptomic analysis

We defined a gene set from the overlap of differentially expressed genes from a microarray comparing GF vs. recolonized animals and in a RNA-seq experiment comparing 12°C vs. 18°C rearing temperature (55 genes in total) (29, 52). For technical details of RNA-seq and microarray analysis, see (29, 52). This gene set was associated with ranks according mean expression, p-value and fold change for both conditions (decreasing order for fold change and mean expression, increasing order for p-values). The ranks for each category were summed, to generate a score which was used to sort the genes. The genes Eco1 (contig 18166) and Eco2 (contig 14187) were at position 2 and 7 in this list. We did not consider the first gene in the table (a serine acetyltransferase, contig 18585) as it was predicted to be associated with metabolic changes which was not the focus of this study. Eco2 was considered only after further investigation of Eco1 and appearing also in a high rank in this list.

### In-situ hybridization

Whole open reading frame of *eco1* and *eco2* where cloned into pGEM-T (*Promega*). DIG-labeled probes where generated using T7/SP6 transcriptions start sites and the digoxigenin RNA Labeling Mix kit (*Roche*) after manufactures instructions. Hybridization was performed as described previously (89), in short: animals were relaxed with 2 % urethan for 2 min at 4 °C, fixed in 4 % PFA over night at 4 °C, washed three times á 10 min with PBT (PBS + 0.1 % Tween), and bleached in methanol (100 %) three times for minimum 30 min. Methanol was washed out with four decreasing concentrations of ethanol in PBT (100 %, 75 %, 50 %, 25 % v/v) followed by three times PBT washing steps, all 10 min or longer on a shaker. Samples were digested with 10 mg ml^-1^ Proteinase K in PBT for 20 min at room temperature and digestion was stopped by two subsequent washing steps with 4 mg ml^-1^ glycine in PBT, followed by three PBT washing steps each 10 min on a shaker. In order to acetylate positve charges in the samples and reduce background we treated samples twice with triethanolamine (0.1 M, pH 7.8) 5 min, once more with triethanolamine adding 2.5 µl µm^-1^ acetic anhydrid for 5 min to add another 2.5 µl ml^-1^ acetic anhydrid for another 5 min. The samples where washed three times in PBT, refixed in 4 % PFA over night at 4 °C and washed another three times in PBT to remove fixative. Samples were treated twice with 2x SCC (0.3 M NaCl, 30 mM tri-sodium citrate, pH 7) for 10 min at room temperature and twice for 20 min at 70 °C, to inactivate endogenous phosphatases. To transfer the samples into hybridization solution (5x SSC + 50 % formamide (v/v), 0.02 % Ficoll (w/v), 0.02 % bovine serum albumin (w/v), 0.02 % polyvinylpyrolidone (w/v), 0.1 % Tween20 (v/v), 0.1 % CHAPS (w/v), 100 µg ml^-1^ Heparin), the medium was exchanged to prewarmed 50 % hybridization solution in 2x SSC for 10 min at 57 °C, followed by 10 min in hybridization solution at 57 °C and 2 h blocking in hybridization solution + 100 µg ml^-1^ tRNA at 57 °C. Around 20 ng µl^-1^ *in-situ* probes were denatured for 10 min at 70 °C in 10x SSC + 50 % formamid. The denatured probe was added to the hybridization solution + tRNA to a final concentration of 2 ng µl^-1^ and incubated over night at 57 °C. Sense probes were used equally and served as negative control. To remove the probes and prepare antibody staining, the samples were washed in four steps of decreasing hybridization solution in 2x SSC (100 %, 75 %, 50 %, 25 %) for 10 min at 57 °C each. Afterwards, samples were washed twice in 2x SSC containing 0.1 % CHAPS for 30 min at 57 °C and twice in MAB-T (100 mM maleic acid, 150 mM NaCl, 0.1 % Tween20 (v/v), pH 7.5) for 10 min at room temperature. To block unspecific antibody binding sites, the samples were blocked with MAB-T + 0.1 % bovine serum albumin for 1 h at room temperature and with MAB-T + 0.1 % bovine serum albumin, 20 % heat inactivated sheep serum for 2 h at 4 °C. Anti-digoxigenin alkaline phosphate linked fab fragments (*Roche*) were used to detect probes by incubating the samples with a 1:2000 solution in MAB-T + 0.1 % bovine serum albumin, 20 % heat inactivated sheep serum over night at 4 °C. Excess antibody solution was removed by 8 washes with MAB-T for 15 min at room temperature. Sample staining was prepared by one NTMT (100 mM NaCl, 100 mM TRIS-HCl, 50 mM MgCl2, 0.1 % Tween20 (v/v), pH 9.5) wash and incubation with NTMT + 1 mM Levamisol for 5 min to inhibit unspecific endogenous alkaline phosphatases. Staining was performed by incubation of the samples in NTMT + 2 % NBT/BCIP in dark until a clear staining was obtained. The reaction was stopped with three washes of a*qua dest*. before dehydration in with three steps increasing ethanol concentrations (50 %, 70 %, 100 % in water) was performed and samples were embedded on glas slides in Euparal. Expression profiles of *eco* genes under the different treatments were analyzed under the microscope.

### Immunohistochemistry

Immunostainging was performed using standard procedures (90): animals were relaxed in 2 % urethan for 2 min on ice and fixed in 4 % PFA over night at 4 °C. Fixative was removed with four washes in PBT (PBS + 0.1 % Tween20) for 15 min, cell membranes opened with PBS + 0.5 % TritonX100 for 30 min, and unspecific binding sites blocked with PBT + 1 % bovine serum albumin for 1 h, everything done at room temperature. The primary antibody was a polyclonal rabbit antibody raised against a 14 AA fragment (NSIKENMENFYPVE, AA 52-65, *GeneScript, USA*) of Eco1 and was incubated in PBT + 1 % bovine serum albumin, 10 µg/ml rabbit-α-Eco1-AB over night at 4 °C. The primary antibody was removed with four washes in PBT + 1 % bovine serum albumin for 15 min at room temperature. Secondary anitbody (goat-α-rabbit Alexa Flour 488 coupled, #A11034; LOT 1670152, *Life Technologies*) was incubated at 1 µg/ml in PBT + 1 % bovine serum albumin and incubated for 2 h at room temperature, from this step on samples were kept in dark. Secondary antibody was removed by four washes in PBS + 0.5 % Tween20, 1 % bovine serum albumin for 15 min at room temperature. Actin cytoskeleton staining was performed with rhodamine-phalloidin (#P1951, *Sigma*) in PBT for 1 h at room temperature and afterwards removed with four washes of PBS + 0.5 % Tween20, 1 % bovine serum albumin. To stain the nucleus we incubated the samples with TO-PRO3 iodide AlexaFluor633 (#T3605, LOT 23929W, *Invitrogen*) in PBS + 0.5 % Tween20, 1 % bovine serum albumin for 10 min at room temperature. Afterwards samples were transferred onto glas slides, embedded in Moviol and stored at 4 °C until analysis using a confocal laser scanning microscope (TCS SP1, *Leica*)

### Transplantation experiments

Rings of tissue right beneath the tentacle ring were excised from donor animals and grafted into the ∼1/3 of the body axis from head to foot of the acceptor animal, using fishing strings and polypropylene tubes for fixation of the tissues. Grafts were grown together after 2-3 h and fishing strings were removed. The induction of heads was assessed 2-3 days post transplantation using the fluorescent markers and a binocular. Every form of head structure was evaluated as head, including a single tentacle, a hypostome or a complete hypostome with an tentacle ring.

## Author contributions

J. Taubenheim, S. Franzenburg, T. C. G. Bosch and S. Fraune. designed the experiments; J. Taubenheim, M. Knop, D. Willoweit-Ohl, S. Franzenburg, and J. He conducted experiments. J. Taubenheim, M. Knop and S. Fraune performed data analysis. J. Taubenheim, T. C. G. Bosch and S. Fraune wrote the paper, all authors corrected final version of the paper.

## Acknowledgement

We greatly appreciate provision and microinjection of the *Hydra* embryos by Jörg Wittlieb. We further thank Alexander Klimovich for help in the design of the antibodies and help with the confocal microscopy. This work was funded by the German Research Foundation (DFG, CRC1182). T. C. G. B. gratefully appreciates support from the Canadian Institute for Advanced Research (CIFAR).

## Supplements

**Supplementary Figure 1:**
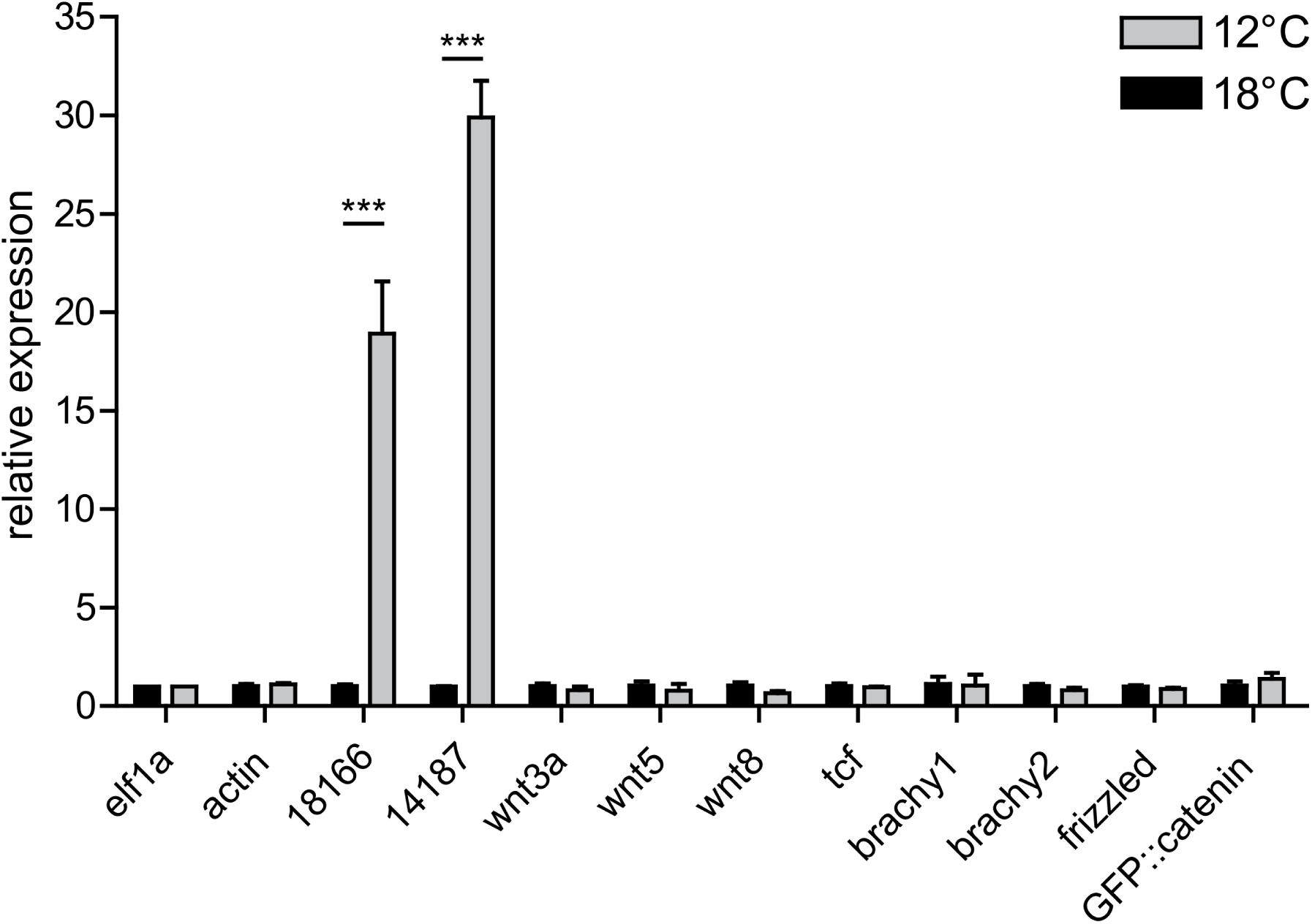
The temperature shift does not change the expression of Wnt pathway components. Temperature change has no impact on the expression of Wnt3a, Wnt5, Wnt8, TCF, Frizzled, head formation associated hox genes (brachyury 1 and 2) or the expression of the β-catenin OE construct, but is associated with an 20 to 30 fold up regulation of the eco genes (n =3, Two-way ANOVA, Bonferroni posttests, *** p<0.001).

**Supplementary Figure 2:**
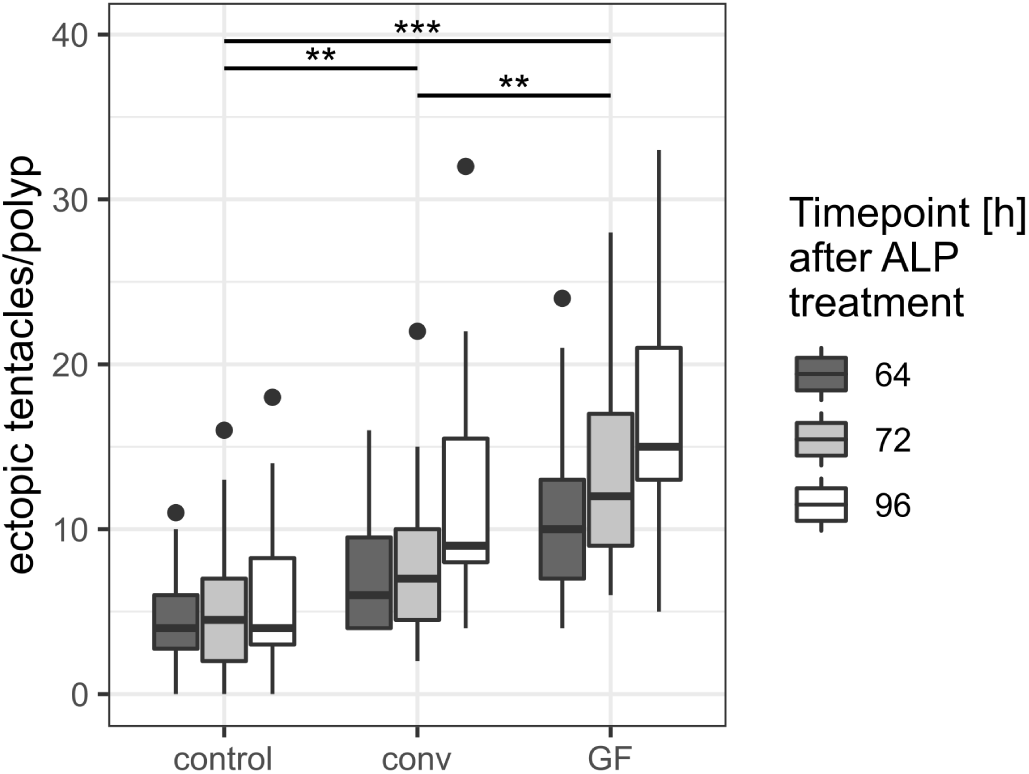
Ectopic tentacle formation after ALP is dependent on the bacterial colonization in *Hydra*. Treatment of germ-free *Hydra* polyps with ALP leads to an increase of ectopic tentacle formation, compared to control and conventionalized animals (recolonized for 4 days prior treatment). The conventionalized and the germ-free animals show some dynamics in tentacle formation, with most tentacles formed 96 h after ALP treatment. Conventionalized *Hydra* represent an intermediate state for tentacle formation after ALP treatment, but is able to mitigate the effect in germ-free animals. n = 15 for conv and GF, 20 for control, fitted generalized linear model with poisson log link function for each time point individually. Multiple comparison for treatment factor with generalized linear hypothesis testing (R package multcomp) and p-values corrected with Benjamini-Hochberg procedure. Asterisks code f or the highest p-value in the comparisons at the different timepoints: *** p < 0.001, ** p < 0.01. See Supplementary Table 1 for test statistics.

**Supplementary Figure 3:**
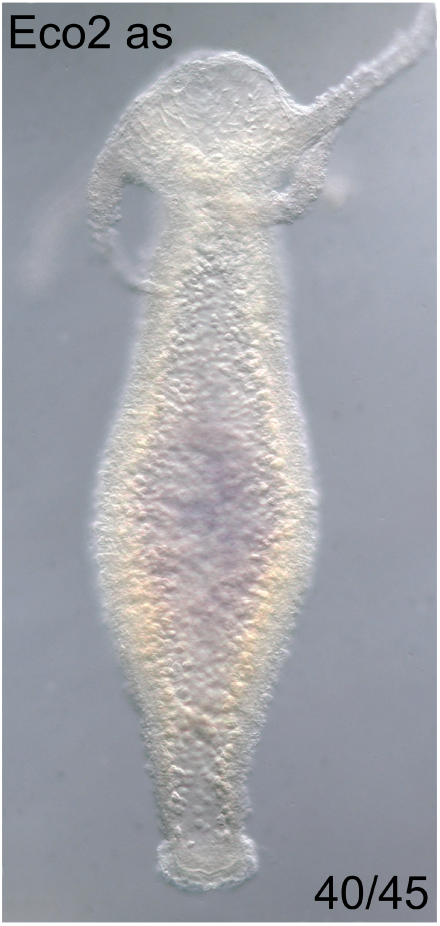
Missing expression signal for *eco2* gene expression after in- situ hybridization. Most of the animals showed no signal after in-situ hybridization with an anti-sense probe. The low fraction of stained animals is caused by the low *eco2* signal at 18°C rearing temperature.

**Supplementary Figure 4:**
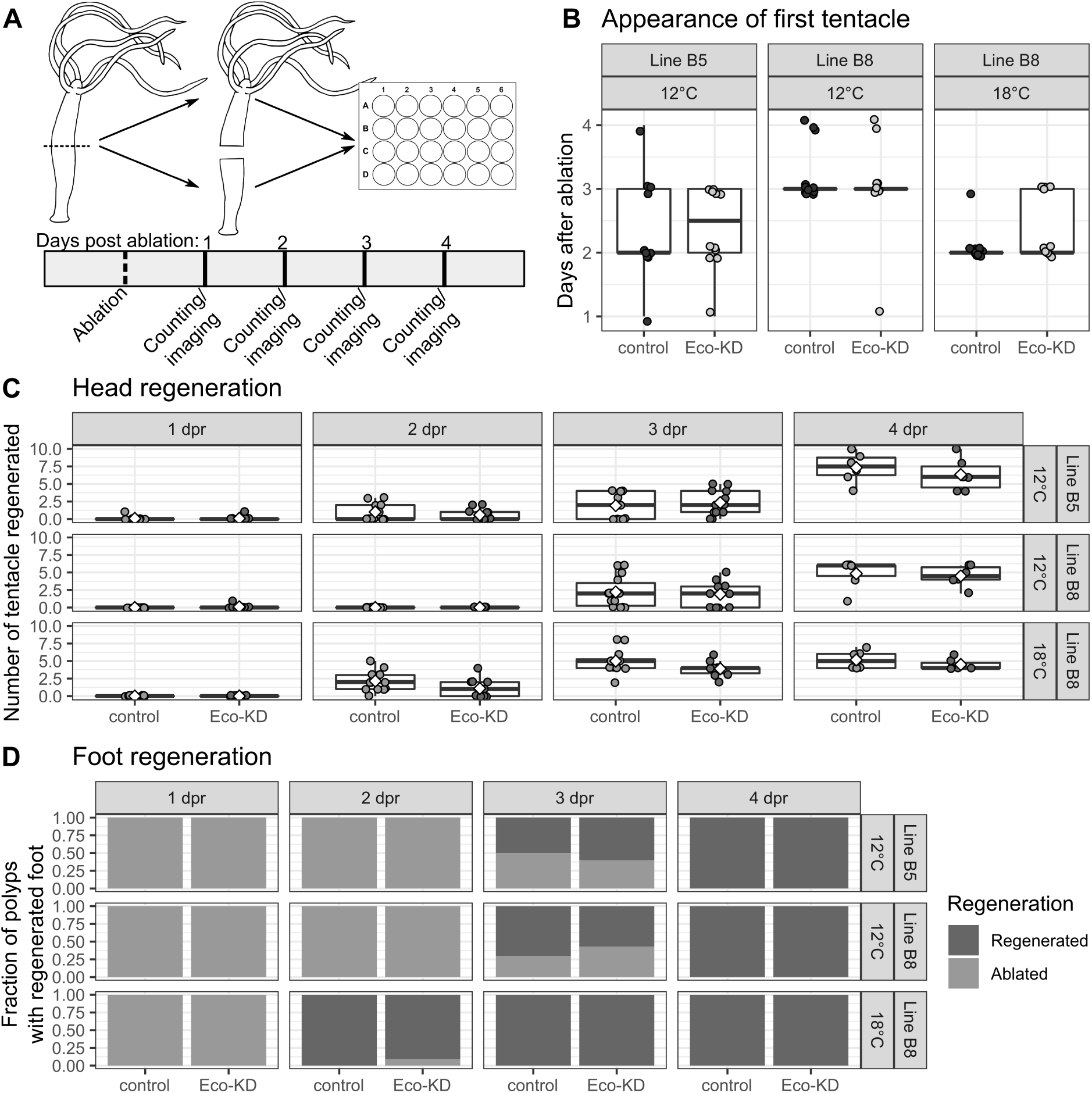
Eco peptides are not involved in the regeneration process of *Hydra*. (A) To test for a role of *eco* genes in pattern formation during regeneration, animals were, randomized, cut in half and each piece was placed individually in a 24-well plate. The regenerating pieces were observed for 3-4 days, counting emerging tentacles and foots every day. (B) The timing of the head regeneration (first appearance of a tentacle) was not altered by the knock-down of the *eco* genes in 12°C as well as 18°C (Wilcoxon-test, with Benjamini-Hochberg correction, n = 9-13 polyps for each group). (C) Number of tentacles at different time points of regeneration was indifferent between Eco-KD and control animals (t- test @dpa 3 and 4, with Benjamini-Hochberg correction, n=6-14 polyps for each group). (D) The fraction of polyps regenerating a foot after ablation was not changed in an Eco-KD background (Fisher’s-exact test @dpa =3 (12°C) or 2 (18°C), n = 6-14 polyps for each group).

**Supplementary Figure 5:**
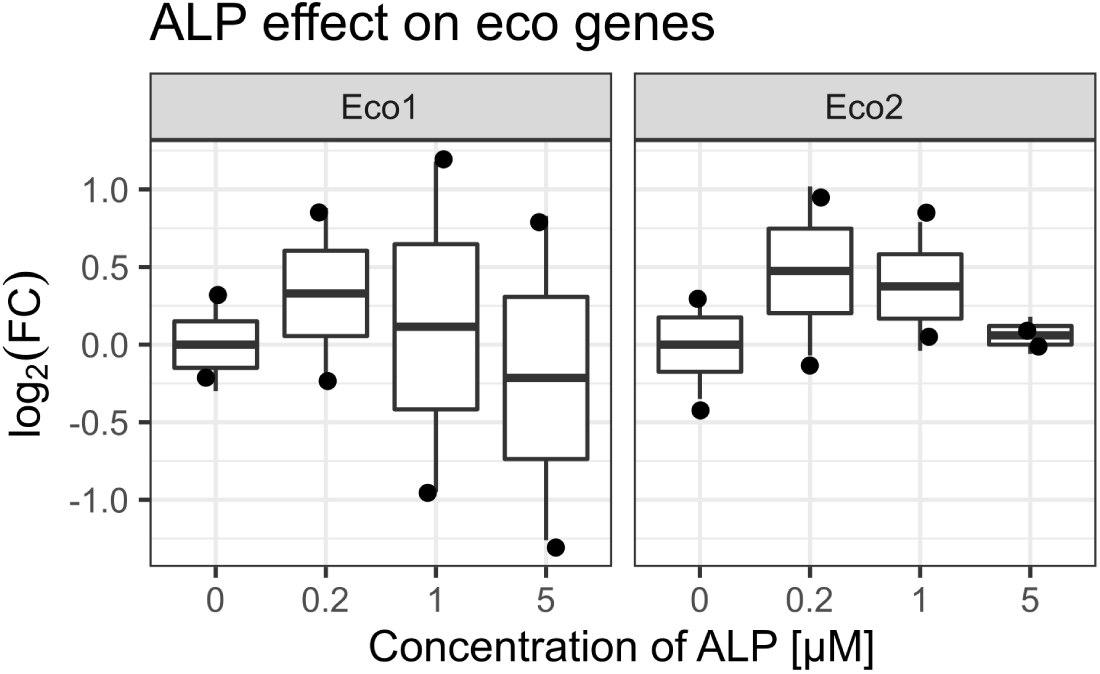
ALP treatment does not result in expression changes of eco genes. Animals treated with different concentrations of ALP for 24 h showed now significant difference in the gene expression for eco genes (ANOVA, gene-stratified p = 0.532, see Supplementary Table 4).

## References

1. McGlashan JK, Spencer R-J, Old JM (2012) Embryonic communication in the nest: metabolic responses of reptilian embryos to developmental rates of siblings. Proc R Soc B Biol Sci 279(1734):1709–1715.

2. Callier V, Nijhout HF (2011) Control of body size by oxygen supply reveals size-dependent and size-independent mechanisms of molting and metamorphosis. Proc Natl Acad Sci 108(35):14664–14669.

3. Zera AJ, Tiebel KC (1988) Brachypterizing effect of group rearing, juvenile hormone III and methoprene in the wing-dimorphic cricket, Gryllus rubens. J Insect Physiol 34(6):489–498.

4. Ogawa A, Streit A, Antebi A, Sommer RJ (2009) A Conserved Endocrine Mechanism Controls the Formation of Dauer and Infective Larvae in Nematodes. Curr Biol 19(1):67–71.

5. Shikuma NJ, Antoshechkin I, Medeiros JM, Pilhofer M, Newman DK (2016) Stepwise metamorphosis of the tubeworm Hydroides elegans is mediated by a bacterial inducer and MAPK signaling. Proc Natl Acad Sci 113(36):10097–10102.

6. West-Eberhard MJ (2003) Developmental plasticity and evolution (Oxford University Press) Available at: https://books.google.de/books?id=iBkQyA2PkxEC&dq=Developmental+Plasticity+and+Evolution&hl=de&source=gbs_navlinks_s [Accessed April 25, 2019].

7. Gilbert SF, Bosch TCG, Ledón-Rettig C (2015) Eco-Evo-Devo: Developmental symbiosis and developmental plasticity as evolutionary agents. Nat Rev Genet 16(10):611–622.

8. Abouheif E, Favé MJ, Ibarrarán-Viniegra AS, Lesoway MP, Rafiqi AM, Rajakumar R (2014) Eco-Evo-Devo: The time has come. Adv Exp Med Biol 781:107–125.

9. Ghosh SM, Testa ND, Shingleton AW (2013) Temperature-size rule is mediated by thermal plasticity of critical size in Drosophila melanogaster. Proc R Soc B Biol Sci 280(1760):20130174–20130174.

10. Atkinson D (1994) Temperature and Organism Size—A Biological Law for Ectotherms?, pp 1–58.

11. Gillooly JF, Charnov EL, West GB, Savage VM, Brown JH (2002) Effects of size and temperature on developmental time. Nature 417(6884):70–73.

12. Bergmann C (1848) Über die Verhältnisse der Wärmeökonomie der Thiere zu ihrer Grösse. Available at: https://books.google.de/books?hl=de&lr=&id=EHo-AAAAcAAJ&oi=fnd&pg=PA3&dq=Bergmann+1848&ots=YkWTtFnaf5&sig=5wW6LvN-JR7V7g5L_Ewjhh8Fjws [Accessed April 29, 2019].

13. Li Q, Gong Z (2015) Cold-sensing regulates Drosophila growth through insulin-producing cells. Nat Commun 6:1–11.

14. Fraune S, Bosch TC (2010) Why bacteria matter in animal development and evolution. Bioessays 32(7):571–580.

15. Landmann F, Foster JM, Michalski ML, Slatko BE, Sullivan W (2014) Co-evolution between an Endosymbiont and Its Nematode Host: Wolbachia Asymmetric Posterior Localization and AP Polarity Establishment. PLoS Negl Trop Dis 8(8):e3096.

16. Landmann F, Voronin D, Sullivan W, Taylor MJ (2011) Anti-filarial Activity of Antibiotic Therapy Is Due to Extensive Apoptosis after Wolbachia Depletion from Filarial Nematodes. PLoS Pathog 7(11):e1002351.

17. Montgomery MK, McFall-Ngai M (1994) Bacterial Symbionts Induce Host Organ Morphogenesis during Early Postembryonic Development of the Squid Euprymna Scolopes. Development 120(7):1719–1729.

18. De Vadder F, Grasset E, Holm LM, Karsenty G, Macpherson AJ, Olofsson LE, Bäckhed F (2018) Gut microbiota regulates maturation of the adult enteric nervous system via enteric serotonin networks. Proc Natl Acad Sci U S A 115(25):6458–6463.

19. Hedgecock EM, Russell RL (1975) Normal and mutant thermotaxis in the nematode Caenorhabditis elegans. Proc Natl Acad Sci 72(10):4061–4065.

20. Kammenga JE, Doroszuk A, Riksen JAG, Hazendonk E, Spiridon L, Petrescu A-J, Tijsterman M, Plasterk RHA, Bakker J (2007) A Caenorhabditis elegans wild type defies the temperature-size rule owing to a single nucleotide polymorphism in tra-3. PLoS Genet 3(3):e34.

21. Okamoto N, Nishimura T (2015) Signaling from Glia and Cholinergic Neurons Controls Nutrient-Dependent Production of an Insulin-like Peptide for Drosophila Body Growth. Dev Cell 35(3):295–310.

22. Fujiwara M, Sengupta P, McIntire SL (2002) Regulation of body size and behavioral state of C. elegans by sensory perception and the EGL-4 cGMP-dependent protein kinase. Neuron 36(6):1091–102.

23. Sawala A, Gould AP (2017) The sex of specific neurons controls female body growth in Drosophila. PLOS Biol 15(10):e2002252.

24. Pietschke C, Treitz C, Forêt S, Schultze A, Künzel S, Tholey A, Bosch TCG, Fraune S (2017) Host modification of a bacterial quorum-sensing signal induces a phenotypic switch in bacterial symbionts. Proc Natl Acad Sci U S A 114(40). doi: 10.1073/pnas.1706879114.

25. Fraune S, Anton-Erxleben F, Augustin R, Franzenburg S, Knop M, Schröder K, Willoweit-Ohl D, Bosch TCG (2015) Bacteria-bacteria interactions within the microbiota of the ancestral metazoan Hydra contribute to fungal resistance. ISME J 9(7). doi: 10.1038/ismej.2014.239.

26. Franzenburg S, Walter J, Künzel S, Wang J, Baines JF, Bosch TCG, Fraune S (2013) Distinct antimicrobial peptide expression determines host species-specific bacterial associations. Proc Natl Acad Sci U S A 110(39). doi: 10.1073/pnas.1304960110.

27. Mortzfeld BM, Taubenheim J, Fraune S, Klimovich AV, Bosch TCG (2018) Stem cell transcription factor FoxO controls microbiome resilience in hydra. Front Microbiol 9(APR). doi: 10.3389/fmicb.2018.00629.

28. Fraune S, Bosch TCG (2007) Long-term maintenance of species-specific bacterial microbiota in the basal metazoan Hydra. Proc Natl Acad Sci U S A 104(32):13146–13151.

29. Mortzfeld BM, Taubenheim J, Klimovich A V., Fraune S, Rosenstiel P, Bosch TCG (2019) Temperature and insulin signaling regulate body size in Hydra by the Wnt and TGF-beta pathways. Nat Commun 10(1):3257.

30. Bosch TCG (2007) Why polyps regenerate and we don’t: Towards a cellular and molecular framework for Hydra regeneration. Dev Biol 303(2):421–433.

31. Field KG, Olsen GJ, Lane DJ, Giovannoni SJ, Ghiselin MT, Raff EC, Pace NR, Raff RA (1988) Molecular Phylogeny of the Animal Kingdom. Science (80-) 239(2):748–753.

32. Campbell RD (1967) Tissue dynamics of steady state growth in Hydra littoralis. Dev Biol 15(5):487–502.

33. Campbell RD (1967) Tissue dynamics of steady state growth inHydra littoralis. II. Patterns of tissue movement. J Morphol 121(1):19–28.

34. Campbell RD (1967) Tissue dynamics of steady state growth inhydra littoralis. III. Behavior of specific cell types during tissue movements. J Exp Zool 164(3):379–391.

35. Holstein TW, Hobmayer E, David CN (1991) Pattern of epithelial cell cycling in hydra. Dev Biol 148(2):602–611.

36. Hobmayer B, Rentzsch F, Kuhn K, Happel CM, von Laue CC, Snyder P, Rothbächer U, Holstein TW (2000) WNT signalling molecules act in axis formation in the diploblastic metazoan Hydra. Nature 407(6801):186–9.

37. Lengfeld T, Watanabe H, Simakov O, Lindgens D, Gee L, Law L, Schmidt HA, Özbek S, Bode H, Holstein TW (2009) Multiple Wnts are involved in Hydra organizer formation and regeneration. Dev Biol 330(1):186–199.

38. Gee L, Hartig J, Law L, Wittlieb J, Khalturin K, Bosch TCG, Bode HR (2010) beta-catenin plays a central role in setting up the head organizer in hydra. Dev Biol 340(1):116–24.

39. MacWilliams HK (1983) Hydra transplantation phenomena and the mechanism of Hydra head regeneration. Dev Biol 96(1):217–238.

40. Browne EN (1909) The production of new hydranths in Hydra by the insertion of small grafts. J Exp Zool 7(1):1–23.

41. Turing A (1952) The chemical basis of morphogenesis. Philos Trans R Soc Lond B Biol Sci 237(641):37–72.

42. Gierer A, Meinhardt H (1972) A theory of biological pattern formation. Kybernetik 12(1):30–39.

43. Broun M, Gee L, Reinhardt B, Bode HR (2005) Formation of the head organizer in hydra involves the canonical Wnt pathway. Development 132(12):2907–2916.

44. Klimovich A, Rehm A, Wittlieb J, Herbst E-M, Benavente R, Bosch TCG (2018) Non-senescent Hydra tolerates severe disturbances in the nuclear lamina. Aging (Albany NY) 10(5):951–972.

45. Bosch TCG, Adamska M, Augustin R, Domazet-Loso T, Foret S, Fraune S, Funayama N, Grasis J, Hamada M, Hatta M, et al. (2014) How do environmental factors influence life cycles and development? An experimental framework for early-diverging metazoans. Bioessays 36(12):1185–94.

46. Boehm A-M, Rosenstiel P, Bosch TCG (2013) Stem cells and aging from a quasi-immortal point of view. BioEssays 35(11):994–1003.

47. Galliot B, Buzgariu W, Schenkelaars Q, Wenger Y (2018) Non-developmental dimensions of adult regeneration in Hydra. Int J Dev Biol 62(6-7–8):373–381.

48. Gufler S, Artes B, Bielen H, Krainer I, Eder M-K, Falschlunger J, Bollmann A, Ostermann T, Valovka T, Hartl M, et al. (2018) β-Catenin acts in a position-independent regeneration response in the simple eumetazoan Hydra. Dev Biol 433(2):310–323.

49. Bisbee JW (1973) Size determination in Hydra: The roles of growth and budding. Development 30(1).

50. Otto JJ, Campbell RD (1977) Tissue economics of hydra: regulation of cell cycle, animal size and development by controlled feeding rates. J Cell Sci 28(1).

51. Leost M, Schultz C, Link A, Wu YZ, Biernat J, Mandelkow EM, Bibb JA, Snyder GL, Greengard P, Zaharevitz DW, et al. (2000) Paullones are potent inhibitors of glycogen synthase kinase-3beta and cyclin-dependent kinase 5/p25. Eur J Biochem 267(19):5983–94.

52. Franzenburg S, Fraune S, Kunzel S, Baines JF, Domazet-Loso T, Bosch TCG (2012) MyD88-deficient Hydra reveal an ancient function of TLR signaling in sensing bacterial colonizers. Proc Natl Acad Sci 109(47):19374–19379.

53. Khalturin K, Hemmrich G, Fraune S, Augustin R, Bosch TCG (2009) More than just orphans: are taxonomically-restricted genes important in evolution? Trends Genet 25(9):404–413.

54. Tautz D, Domazet-Lošo T (2011) The evolutionary origin of orphan genes. Nat Rev Genet 12(10):692–702.

55. Turing AM (1952) The Chemical Basis of Morphogenesis. Philos Trans R Soc B Biol Sci 237(641):37–72.

56. Meinhardt H, Gierer A (1974) Applications of a theory of biological pattern formation based on lateral inhibition. J Cell Sci 15(2):321–46.

57. Greber MJ, David CN, Holstein TW (1992) A quantitative method for separation of living Hydra cells. Roux’s Arch Dev Biol 201(5):296–300.

58. MacWilliams HK (1983) Hydra transplantation phenomena and the mechanism of Hydra head regeneration. II. Properties of the head activation. Dev Biol 96(1):239–57.

59. MacWilliams HK (1983) Hydra transplantation phenomena and the mechanism of Hydra head regeneration. I. Properties of the head inhibition. Dev Biol 96(1):217–238.

60. Takano J, Sugiyama T (1983) Genetic analysis of developmental mechanisms in hydra. VIII. Head-activation and head-inhibition potentials of a slow-budding strain (L4). J Embryol Exp Morphol 78(2):141–68.

61. Holstein TW (2012) The evolution of the Wnt pathway. Cold Spring Harb Perspect Biol 4(7):a007922.

62. Watanabe H, Schmidt HA, Kuhn A, Höger SK, Kocagöz Y, Laumann-Lipp N, Özbek S, Holstein TW (2014) Nodal signalling determines biradial asymmetry in Hydra. Nature 515(7525):112–115.

63. Khalturin K, Anton-Erxleben F, Milde S, Plötz C, Wittlieb J, Hemmrich G, Bosch TCG (2007) Transgenic stem cells in Hydra reveal an early evolutionary origin for key elements controlling self-renewal and differentiation. Dev Biol 309(1):32–44.

64. Hartl M, Glasauer S, Gufler S, Raffeiner A, Puglisi K, Breuker K, Bister K, Hobmayer B (2019) Differential regulation of myc homologs by Wnt/β-Catenin signaling in the early metazoan Hydra. FEBS J 286(12):2295–2310.

65. Guder C, Pinho S, Nacak TG, Schmidt HA, Hobmayer B, Niehrs C, Holstein TW (2006) An ancient Wnt-dickkopf antagonism in Hydra. Development 133(5):901–911.

66. Augustin R, Franke A, Khalturin K, Kiko R, Siebert S, Hemmrich G, Bosch TCG (2006) Dickkopf related genes are components of the positional value gradient in Hydra. Dev Biol 296(1):62–70.

67. Vogg MC, Beccari L, Iglesias Ollé L, Rampon C, Vriz S, Perruchoud C, Wenger Y, Galliot B (2019) An evolutionarily-conserved Wnt3/β-catenin/Sp5 feedback loop restricts head organizer activity in Hydra. Nat Commun 10(1):312.

68. Lommel M, Strompen J, Hellewell AL, Balasubramanian GP, Christofidou ED, Thomson AR, Boyle AL, Woolfson DN, Puglisi K, Hartl M, et al. (2018) Hydra Mesoglea Proteome Identifies Thrombospondin as a Conserved Component Active in Head Organizer Restriction. Sci Rep 8(1):1–18.

69. Reddy PC, Gungi A, Ubhe S, Pradhan SJ, Kolte A, Galande S (2019) Molecular signature of an ancient organizer regulated by Wnt/β-catenin signalling during primary body axis patterning in Hydra. Commun Biol 2(1):1–11.

70. Cardenas M, Fabila Y V., Yum S, Cerbon J, Böhmer FD, Wetzker R, Fujisawa T, Bosch TCG, Salgado LM (2000) Selective protein kinase inhibitors block head-specific differentiation in hydra. Cell Signal 12(9–10):649–658.

71. Fabila Y, Navarro L, Fujisawa T, Bode HR, Salgado LM (2002) Selective inhibition of protein kinases blocks the formation of a new axis, the beginning of budding, in Hydra. Mech Dev 119(2):157–164.

72. Schüler A, Schmitz G, Reft A, Özbek S, Thurm U, Bornberg-Bauer E (2015) The Rise and Fall of TRP-N, an Ancient Family of Mechanogated Ion Channels, in Metazoa. Genome Biol Evol 7(6):1713–1727.

73. Malafoglia V, Traversetti L, Del Grosso F, Scalici M, Lauro F, Russo V, Persichini T, Salvemini D, Mollace V, Fini M, et al. (2016) Transient Receptor Potential Melastatin-3 (TRPM3) Mediates Nociceptive-Like Responses in Hydra vulgaris. PLoS One 11(3):e0151386.

74. Cheesman SE, Neal JT, Mittge E, Seredick BM, Guillemin K (2011) Epithelial cell proliferation in the developing zebrafish intestine is regulated by the Wnt pathway and microbial signaling via Myd88. Proc Natl Acad Sci 108(Supplement_1):4570–4577.

75. Franco AT, Israel DA, Washington MK, Krishna U, Fox JG, Rogers AB, Neish AS, Collier-Hyams L, Perez-Perez GI, Hatakeyama M, et al. (2005) Activation of β-catenin by carcinogenic Helicobacter pylori. Proc Natl Acad Sci U S A 102(30):10646–10651.

76. Murata-Kamiya N, Kurashima Y, Teishikata Y, Yamahashi Y, Saito Y, Higashi H, Aburatani H, Akiyama T, Peek RM, Azuma T, et al. (2007) Helicobacter pylori CagA interacts with E-cadherin and deregulates the β-catenin signal that promotes intestinal transdifferentiation in gastric epithelial cells. Oncogene 26(32):4617–4626.

77. Heath-Pagliuso S, Rogers WJ, Tullis K, Seidel SD, Cenijn PH, Brouwer A, Denison MS (1998) Activation of the Ah Receptor by Tryptophan and Tryptophan Metabolites †. Biochemistry 37(33):11508–11515.

78. Kawajiri K, Kobayashi Y, Ohtake F, Ikuta T, Matsushima Y, Mimura J, Pettersson S, Pollenz RS, Sakaki T, Hirokawa T, et al. (2009) Aryl hydrocarbon receptor suppresses intestinal carcinogenesis in ApcMin/+ mice with natural ligands. Proc Natl Acad Sci 106(32):13481–13486.

79. Li L, Zheng W, Zhu Y, Ye H, Tang B, Arendsee ZW, Jones D, Li R, Ortiz D, Zhao X, et al. (2015) QQS orphan gene regulates carbon and nitrogen partitioning across species via NF-YC interactions. Proc Natl Acad Sci 112(47):14734–14739.

80. Li L, Foster CM, Gan Q, Nettleton D, James MG, Myers AM, Wurtele ES (2009) Identification of the novel protein QQS as a component of the starch metabolic network in Arabidopsis leaves. Plant J 58(3):485–498.

81. Santos ME, Le Bouquin A, Crumière AJJ, Khila A (2017) Taxon-restricted genes at the origin of a novel trait allowing access to a new environment. Science (80-) 358(6361):386–390.

82. Khalturin K, Anton-Erxleben F, Sassmann S, Wittlieb J, Hemmrich G, Bosch TC (2008) A novel gene family controls species-specific morphological traits in Hydra. PLoS Biol 6(11):e278.

83. Sackton TB, Lazzaro BP, Schlenke TA, Evans JD, Hultmark D, Clark AG (2007) Dynamic evolution of the innate immune system in Drosophila. Nat Genet 39(12):1461–1468.

84. Colbourne JK, Pfrender ME, Gilbert D, Thomas WK, Tucker A, Oakley TH, Tokishita S, Aerts A, Arnold GJ, Basu MK, et al. (2011) The Ecoresponsive Genome of Daphnia pulex. Science (80-) 331(6017):555–561.

85. Kuo C-H, Kissinger JC (2008) Consistent and contrasting properties of lineage-specific genes in the apicomplexan parasites Plasmodium and Theileria. BMC Evol Biol 8(1):108.

86. Schwentner M, Bosch TCG (2015) Revisiting the age, evolutionary history and species level diversity of the genus Hydra (Cnidaria: Hydrozoa). Mol Phylogenet Evol 91:41–55.

87. Hemmrich G, Anokhin B, Zacharias H, Bosch TCG (2007) Molecular phylogenetics in Hydra, a classical model in evolutionary developmental biology. Mol Phylogenet Evol 44(1):281–290.

88. Pfaffl MW (2001) A new mathematical model for relative quantification in real-time RT-PCR. Nucleic Acids Res 29(9):e45.

89. Grens A, Gee L, Fisher DA, Bode HR (1996) CnNK-2,an NK-2 homeobox gene, has a role in patterning the basal end of the axis in *Hydra*. Dev Biol 180(2):473–488.

90. Engel U, Ozbek S, Streitwolf-Engel R, Petri B, Lottspeich F, Holstein TW (2002) Nowa, a novel protein with minicollagen Cys-rich domains, is involved in nematocyst formation in Hydra. J Cell Sci 115(Pt 20):3923–3934.

